## Supplementary Figures and Legends for "Together, Neuropilin 1 and Neuropilin 2 direct α5 integrin trafficking through GTPase-activating-protein p120RasGAP in endothelial cells to promote fibronectin fibrillogenesis"

**Supplementary Figure legends**

**Suppl. Figure 1:** A) Chord plot showing pathways associated with the top 10 proteins (sorted by intensity score) associating with α5 integrin. B-D) STRING interaction networks identifying associations between α5 integrin and RabGTPase proteins (B), myosin motor proteins (C) and Endocytic trafficking-associated proteins (D).

**Suppl. Figure 2:** A) Densitometric analysis of total expression levels of α5 integrin, αV integrin and paxillin by Western blotting as shown in Figure 2G, N = 3.

**Suppl. Figure 3:** A) Total cell lysate input showing expression levels of tensin-1 by Western blotting in siRNA-depleted ECs. B) Supporting densitometric analysis of A), N = 3.

**Suppl. Figure 4:** A) Recycling assay schematic. Surface proteins were biotin labelled prior to incubation at 37 ◦C to stimulate receptor internalisation. Following MESNA stripping, internalised proteins were allowed to recycle for the indicated timepoints at 37 ◦C alongside a MESNA stripped control. EC lysates were immunoprecipitated against biotin to detect the recycled fraction of total α5 integrin by SDS-PAGE and Western blotting. B-C) Quantification of siRNA-depletion of p120RasGAP and Rab21 from total cell lysates, N = 3, N = 2 respectively.

**Suppl. Figure 5:** A) Densitometric quantification of FAK phosphorylation at $Try^{397}$ and $Tyr^{407}$ residues relative to total FAK or β-actin expression in Ctrl and siRNA-depleted ECs, N = 3. B) Quantification of siRNA-depletion of Rab11 from total cell lysates, N = 4. C) Representative confocal microscopy images showing colocalisation between α5 integrin and EEA1 in Ctrl and *si*Rab11 ECs fixed at 180 minutes. D) Representative confocal microscopy images showing colocalisation between α5 integrin and Rab7 in Ctrl and *si*Rab11 ECs fixed at 180 minutes. E) Representative confocal microscopy images showing $GM130^+$ Golgi-body positioning and F-actin protrusions in Ctrl and *si*Rab11 ECs at the scratch-wound edge. Arrows indicate correctly positioned Golgi-bodies. Asterisks indicate statistical significance.

**Suppl. Figure 6:** A) Representative confocal microscopy images showing $ERG1/2^+$ ECs in Pdgfb-$iCreER^{T2}$-negative, $NRP1^{flfl}.EC^{KO}$, $NRP2^{flfl}.EC^{KO}$ and $NRP1^{flfl}NRP2^{flfl}.EC^{KO}$ retinas harvested at P6. B) Quantification of $ERG1/2^+$ EC density shown as relative percentages of respective littermate control animals, N ≥ 2 (n ≥ 4). C) Representative confocal microscopy images showing BS-1 $lectin^-$ collagen $IV^+$ regressed vessels in Pdgfb-$iCreER^{T2}$ negative and $NRP1^{flfl}NRP2^{flfl}.EC^{KO}$ animals. D) Quantification of raw vessel regression, N = 3 (n ≥ 8). E) Right panel: quantification of filopodial length (shown as relative percentages of respective littermate control animals), and filopodial tortuosity (shown as ratios) in Pdgfb-$iCreER^{T2}$-negative, $NRP1^{flfl}.EC^{KO}$, $NRP2^{flfl}.EC^{KO}$ and $NRP1^{flfl}NRP2^{flfl}.EC^{KO}$ animals, (n = 5). F) Individual polar plots showing Golgi-body orientation in Pdgfb-$iCreER^{T2}$ negative and $NRP1^{flfl}NRP2^{flfl}.EC^{KO}$ animals (n = 50 ECs). Asterixis indicate statistical significance.

(Suppl. Figure 1)

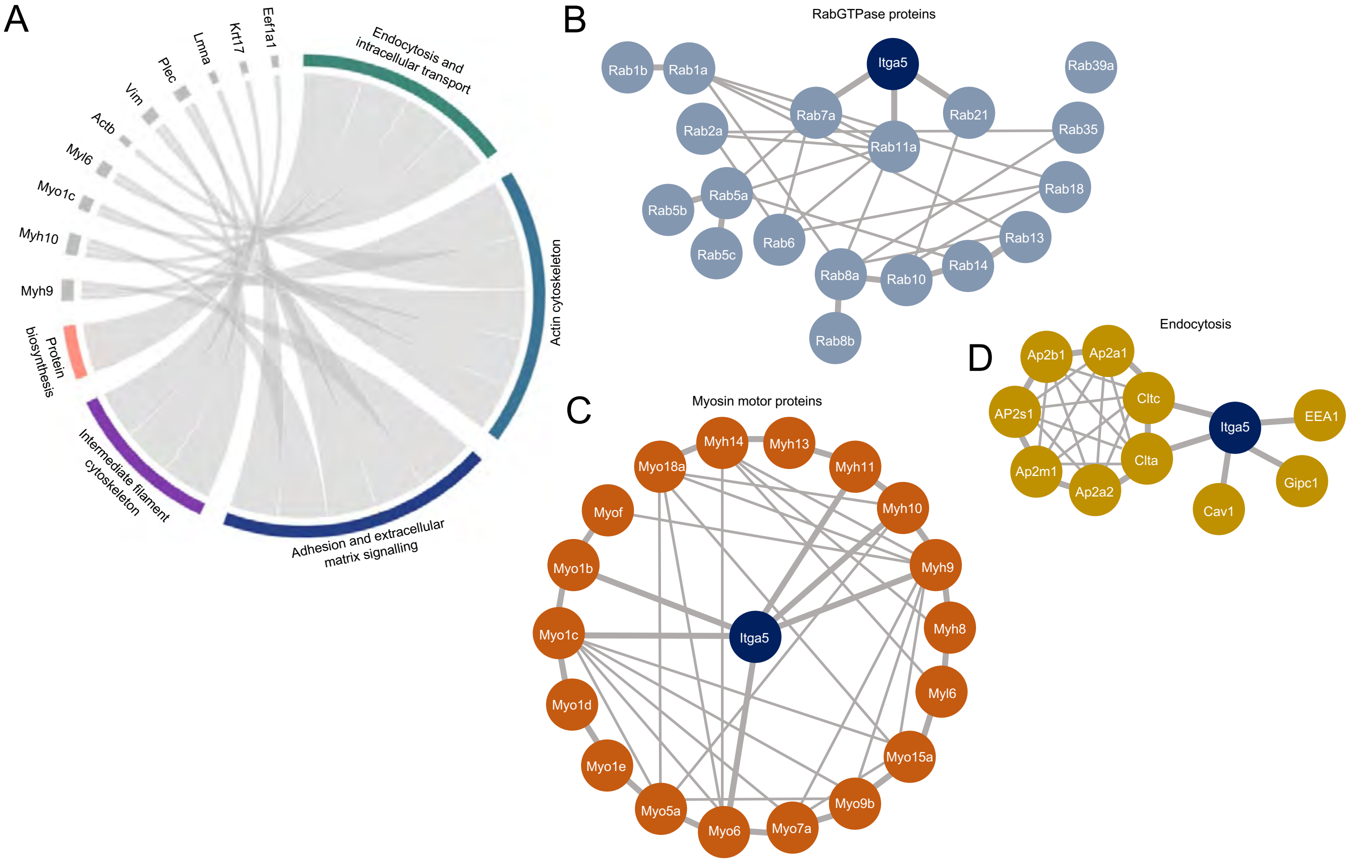

(Suppl. Figure 2)

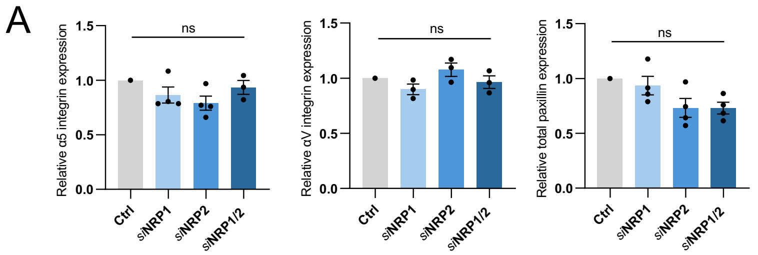


(Suppl. Figure 3)

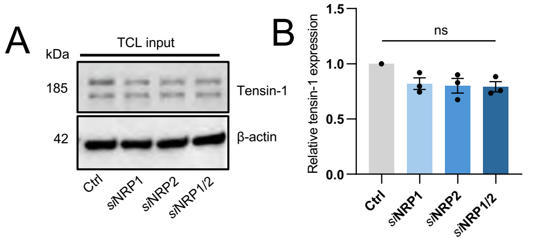


(Suppl. Figure 4)

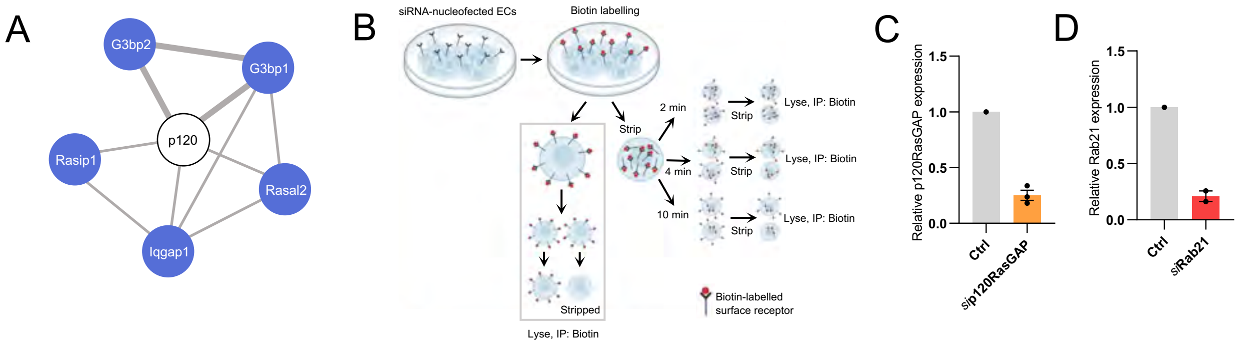


(Suppl. Figure 5)

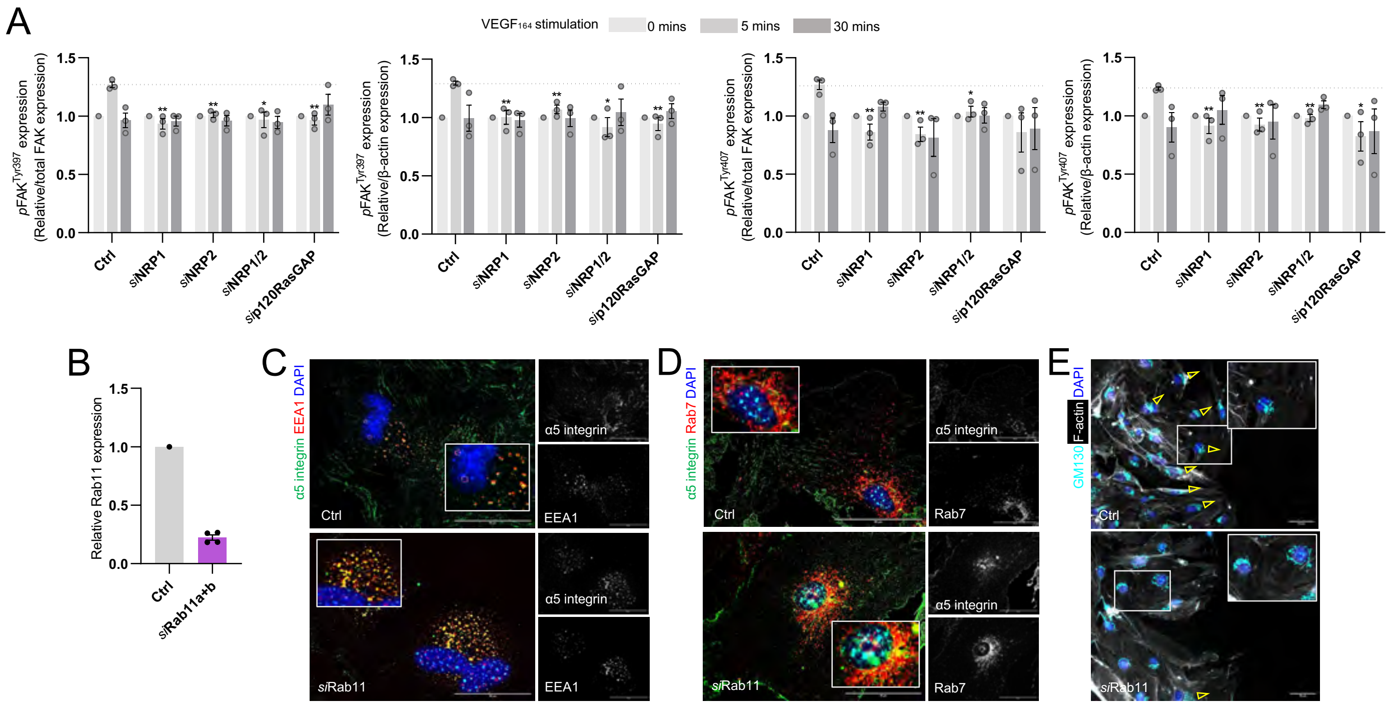

(Suppl. Figure 6)

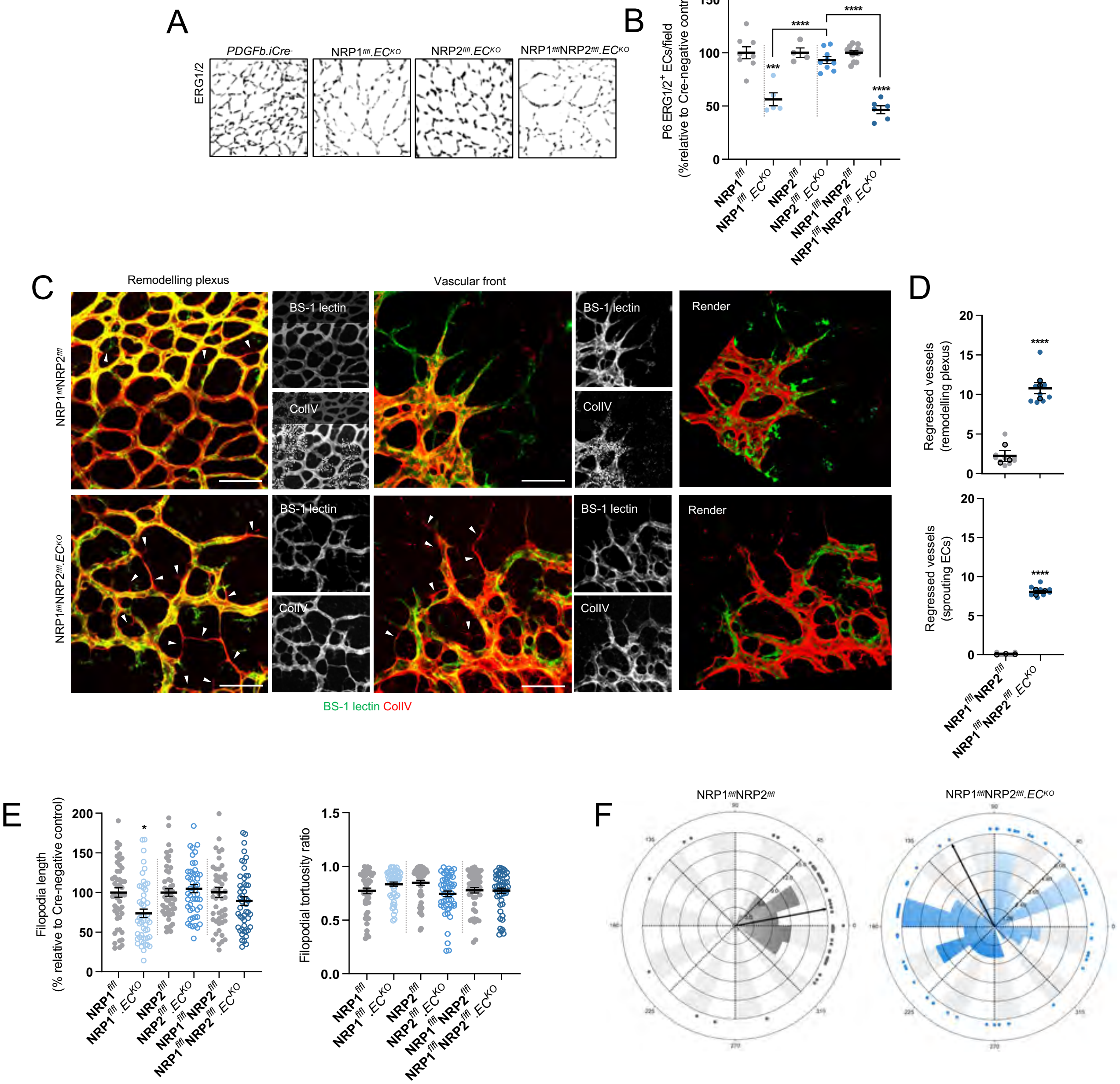
